# Evaluation of a nanopore sequencing strategy on bacterial communities from marine sediments

**DOI:** 10.1101/2023.06.06.541006

**Authors:** Alice Lemoinne, Guillaume Dirberg, Myriam Georges, Tony Robinet

## Abstract

Following the development of high-throughput DNA sequencers, environmental prokaryotic communities were usually described by metabarcoding on short markers of the 16S domain. Among third generation sequencers, that offered the possibility to sequence the full 16s domain, the portable MinION from Oxford Nanopore was undervalued for metabarcoding because of its relatively higher error rate per read. Here we illustrate the limits and benefits of Nanopore sequencing devices by comparing the prokaryotic community structure in a mock community and 52 sediment samples from mangrove sites, inferred from full-length 16S long-reads (16S-FL, *ca*. 1.5 kpb) on a MinION device, with those inferred from partial 16S short-reads (16S- V4V5, *ca*. 0.4kpb, 16S-V4V5) on Illumina MiSeq. 16S-V4V5 and 16S-FL retrieved all the bacterial species from the mock, but Nanopore long-reads overestimated their diversity more than twice. Whether these supplementary OTUs were artefactual or not, they only accounted for *ca*. 10% of the reads. From the sediment samples, with a coverage-based rarefaction of reads and after singletons filtering, Mantel and Procrustean tests of co-inertia showed that bacterial community structures inferred from 16S-V4V5 and 16S- FL were significantly similar, showing both a comparable contrast between sites and a coherent sea-land orientation within sites. In our dataset, 84.7 and 98.8% of the 16S-V4V5 assigned reads were assigned strictly to the same species and genus, respectively, than those detected by 16S-FL. 16S-FL allowed to detect 92.2% of the 309 families and 87.7% of the 448 genera that were detected by the short 16S-V4V5. 16S-FL recorded 973 additional species and 392 genus not detected by 16S-V4V5 (31.5 and 10.4% of the 16S-FL reads, respectively, among which 67.8 and 79.3% were assigned), producted by both primer specificities and diffrent error rates. Thus, our results concluded to an overall similarity between 16S-V4V5 and 16S-FL sequencing strategies for this type of environmental samples.

## 1 INTRODUCTION

The composition and structure of microbial communities are extensively studied through culture- independent methods, based on nucleic acid sequencing, either in bulk DNA extracted from environmental samples (*metagenomic*) or only for DNA markers of interest (gene fragments), amplified from environmental samples before they are sequenced (*metabarcoding*). The metagenomic approach is exempt from amplification bias inherent to metabarcoding (marker specificities, PCR-induced stochasticity) and can produce the useful MAGs (metagenome assembled genomes), but it still faces technical and cost challenges (Taş et al. 2021). The metabarcoding approach remains more widely used, much cheaper, but amplification bias are recurrent : (i) primers choice is crucial and constrained by the maximum size of inserts for second-generation sequencers (400bp for Ion Torrent PGM, 550bp for Illumina MiSeq, Luo et al. 2012) ; (ii) taxa diversity can be overestimated, because of the non-targeted DNA present in the sample (i.e. DNA from the eukaryotic digestive tracts, or extracellular “relic” DNA, Carini et al. 2017) and also because of the ribosomal DNA polymorphism, hidden in individual genomes, an intragenomic variability in the number of duplicates of ribosomal operon, bearing differences in allelic variants between copies (Pereira et al. 2020) ; and (iii) relative abundances of reads per taxa are somehow inaccurate, compared to awaited abundances in the mock samples, a probable consequence of PCR stochasticity and primers specificity.

In high-throughput sequencing (HTS) metabarcoding, the choice of DNA region and primers is known to be crucial for taxa resolution, phylogenetic coverage and sensitivity to community structure. For prokaryotes, none of all the primer pairs that amplifies markers at a convenient size for short-reads (16S-V4V5) strategies (> 550bp for Illumina) can give a complete phylogenetic coverage. Primers spanning over more than one 16S V-region are often preferred, because they improve taxonomic resolution. However, any of these combinations (V1-V2, V3-V4, V4-V5, V6-V8, V7-V9, etc.) showed bias in phylogenetic coverage (Abellan-Schneyder et al. 2021). The 412 bp V4- V5 marker (515F-926R, Parada et al. 2016) covers more broadly the prokaryotic domains (bacteria and archaea), whereas the 438 bp V6-V8 (B969F-BA1406R, Willis et al. 2019) amplifies additional bacterial clades, leading some authors to consider as a best method to combine several short regions along the prokaryotic 16S to minimize these bias (Fuks et al. 2018). However, the multiplication of marker standards for bacteria and archaea also plays against intercomparability.

Third-generation DNA sequencers marked a significant progress for metabarcoding studies, in the fact that the marker size was no longer a technical limitation (tens of kpb for PacBio Sequel II, and no theoretical limit for Nanopore devices), and one can target much more binding sites for primers, improving considerably taxonomic resolution and phylogenetic coverage (Furneaux et al. 2021; Tedersoo et al. 2021; Eshghi Sahraei et al. 2022).

These long-read high-throughput sequencers have been first implemented for sequencing markers from isolated organisms (Schlaeppi et al. 2016; Loit et al. 2019; Maestri et al. 2019). Full-length 16S (16S-FL) environmental metabarcoding has been usually performed on PacBio sequencers, because the Single-Molecule Real-Time (SMRT) technology offers a read quality similar to those of 16S-V4V5 platforms. 16S-FL metabarcoding is mostly used for taxonomic groups in which 16S- V4V5 can lead to a less-accurate assignment, like micro-eukaryotes and specially fungi (Tedersoo et al. 2018; Furneaux et al. 2021; Kolaříková et al. 2021; Eshghi Sahraei et al. 2022; Gueidan and Li 2022), but also a few bacterial phyla (Katiraei et al. 2022). Despite several published works showed the possibility to use Nanopore sequencing for biomedical, environmental or food metabarcoding, by sequencing mock communities of known composition (Benítez-Páez et al. 2016; Davidov et al. 2020; Urban et al. 2021; Toxqui Rodríguez et al. 2023) or by comparing it with an Illumina library sequenced concurrently (J. Shin et al. 2016; H. Shin et al. 2018, Heikema et al. 2020), the great majority of works that we found in literature did not use the Nanopore platform for environmental 16S-FL metabarcoding. This is probably because environmental sample is much more complex than mock communities, and the sequencing depth of Nanopore may be a limiting factor here for large-scale analyses.

Despite raw reads accuracy are similar for PacBio (88-90%) and Nanopore (95-98% on the R9 flow- cells, above 99% for R10.4), the fact that PacBio circular consensus sequence technology (CCS) can align several reads of the same amplicon brings it to an accuracy of >99.9% at 10-fold consensus (Tedersoo et al. 2021). The first 16S-FL third-generation sequencer acknowledged to be suitable for metabarcoding was PacBio Sequel II on fungal complete rRNA operon (ca. 3000 bp, Tedersoo, Tooming-Klunderud and Anslan 2018). Despite its error rate being slightly higher than Illumina, the PacBio 16S-FL sequencing allowed a much better taxonomic resolution, due to the joint powers of ITS1-ITS2 and SSU-LSU flanking regions on the same amplicon.

Promising attempts were made to reach a satisfactory accuracy with Nanopore, by mimicking PacBio with a rolling circle amplification (RCA, Baloğlu et al. 2021) or by flanking, at the two first steps of PCR, each single amplicon with a unique molecular identifier (UMI, Karst et al. 2021). RCA and UMI methods produce a consensus error rate of 0.7% (coverage > 45x) and 0.01% (> 25x) respectively, offering a quality similar to PacBio or Illumina standards (Baloğlu et al. 2021; Karst et al. 2021). The consensus, compared with BLAST (Camacho et al. 2009) to reference sequences of a curated database, could be assigned more accurately to a taxa than standard short markers do (reviewed by Kerkhof 2021). However, lab and downstream bioinformatic workflows are quite complex to implement for ecology scientists, requiring a higher technicity in library preparations and in downstream bioinformatics than for directly sequencing amplicons from environmental samples, as we tested it without success. To date, no environmental metabarcoding based on RCA or UMI protocols have been published.

In community ecology, Nanopore was initially used for simply barcoding individuals with 16S-FL (Maestri et al. 2019), but quickly metabarcoding appeared with Nanopore sequencing alone, to detect pathogen bacterial strains or invasive species, mostly by a metagenomic approach (Brown et al. 2017; Charalampous 2019; Cuscó et al. 2019; Egeter et al. 2022), or to describe eukaryotic communities on the more or less complete rRNA operon (H. Lu et al. 2016; Toxqui Rodríguez et al. 2023). For bacterial communities, studies with a metabarcoding workflow on environmental samples and relying only on Nanopore MinION, aimed at characterizing mouse gut or human respiratory bacteriomes (J. Shin et al. 2016; Ibironke et al. 2020), bacteria associated with algae or plastic debris at sea (H. Shin et al. 2018; Davidov et al. 2020; van der Loos et al. 2021), pathogenic bacteria in food (Planý et al. 2023), fungal communities and biotic interactions (Vass et al. 2022) or pelagic bacteriomes in freshwaters (Urban et al. 2021). In all studies we found, Nanopore was used alone, except for two. Loit et al. (2019) compared it with PacBio CCS for detecting fungal pathogens in plants, concluding that “MinION could be used for rapid and accurate identification of dominant pathogenic organisms and other associated organisms from plant tissues following both amplicon-based and PCR-free metagenomics approaches”. J. Lu et al. (2022) characterized mycobiomes of fungal isolates and environmental samples by sequencing in parallel the full rRNA operon on MinION and the shorter ITS2 on Illumina HiSeq. They concluded that “ITS2 sequencing [was] more biased than full operon sequencing”.

The estimated cost of 1 Gb PacBio sequencing (17€) was lower than Illumina NovaSeq (44€) and MiSeq (56€), but the accessibility to a PacBio sequencer was difficult for this remote place, because of the instrument cost (650 k€ for a PacBio Sequel II) and technicity. Today, the MinION device of Oxford Nanopore Technologies is accessible for 900€, the estimated cost for 1Gb is about 12€, and its smartphone size allows scientists to use it as a field lab device. The portability of the MinIon device is advantageous for molecular ecology scientists located far away from a research center, opening possibilities for studying microbial communities from a field lab, i.e. equipped with usual devices for DNA extraction (mortar, mini-centrifuge, spectrophotometer for DNA drops), PCR (freezer, thermocycler, electrophoresis tank, UV table, ultra-pure water), and libraries making (DNA fluorometer, DNA dryer). Such a field lab is affordable and quite simple to set up for molecular ecologists in remote places or for proposing environmental metabarcoding in the frame of engineering consultancy.

To date, we have found only one published work that has compared Nanopore 16S-FL to Illumina 16S-V4V5 bacterial metabarcoding on the same environmental samples (Heikema et al. 2020). It is noteworthy that Nanopore sequencing introduces more errors than Illumina sequencing, that is more pronounced for artificially generated OTUs using R9 flowcells, even with high accuracy base-calling, compared to R10.4 flowcells that shows a median read accuracy of Q20. Here we propose to evaluate the similarity of bacterial communities described from a mock community and from marine sediments sequenced by 16S 16S-FL on Nanopore MinION device on R9.4 flowcells, and the same samples described on the 16S-V4V5 16S-V4V5 amplified from same DNA extracts on Illumina MiSeq, by addressing a simple question: will these different sequencing strategies conserve (i) the structures of bacterial communities between two neighboring mangrove sites, and (ii) the sea-land orientation of bacterial communities within sites?

## 2 MATERIALS AND METHODS

### 2.1 Sampling sites and sample collection

In June 2019, two sites were selected in the mangrove of Guadeloupe Island, at 6 km of distance each other, for their a priori difference in the level of direct and indirect human pressures (Fig. 1a- b) : the impacted “Rivière salée” site was located on the foreshore of a salty river, close to the city of Pointe-à-Pitre, to its dump and its airport (latitude -61,5469; longitude 16,2594) ; the less-impacted “Babin” site was located in a Ramsar-protected area close to coral reefs (latitude -61,5294 ; longitude 16,3388).

**Figure 1.**
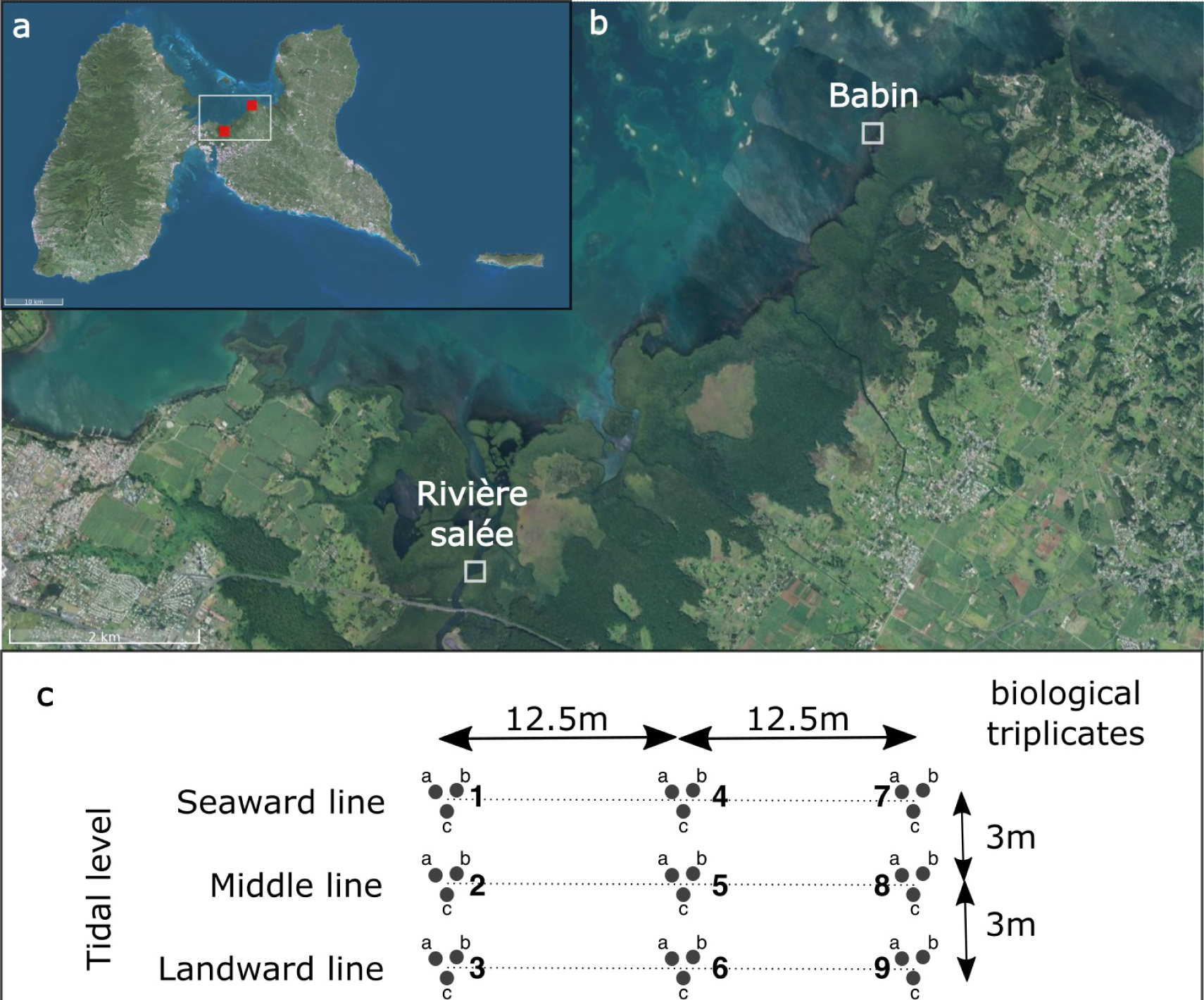
(**a**) Location of sampling sites on Guadeloupe Island (red squares) ; (**b**) zoom on the two sampling sites, with site names ; (**c**) sampling protocol in each site : 3 lines of 3 points, each composed of 3 biological replicates (a, b and c), at 12.5m of distance between each point on each line.

A total of 54 samples of surface sediment were collected on intertidal zone, on 3 lines of 3 points each respectively in each site, each line separated by 3 m to the neighboring line. Points were separated by 12.5 m within a line. Each point was composed of 3 biological replicates (a, b and c), analyzed in the workflow separately (Fig. 1c). The line closest to the sea was the “seaward line”, those closest to the inland mangrove was the “landward line”, the “middle line” was in between. Therefore, each line showed a different time of marine immersion per day. Each replicate was sampled with a sterile syringe and appropriated microbiological precautions, stored in a 50ml Falcon tube, freezed a couple of hours after sampling and preserved at –20° C.

### 2.2 DNA extraction

Samples were sent freezed to metropolitan France, without thawing, then were freeze-dried and crushed to powder in an obsidian mortar, carefully cleaned with an alcoholic tissue between each sample processing. Total genomic DNA from 50mg of dried samples and a standard microbial community (zymoBIOMICS Microbial Community Standard D6300, by ZYMO RESEARCH), here named “*Ze*”, were extracted using the NucleoSpin Soil kit (Macherey-Nagel) with a final elution volume of 50 µl following the manufacturer instructions. After this DNA extraction of samples and Ze, nucleic acid yield and purity were checked using a Nanodrop spectrophotometer (Thermo Fisher Scientific) and the concentration of each sample was equalized to final concentration of 10ng.µl^-1^ on a PCR plate of 96 wells.

### 2.3 Illumina library

In order to limit PCR biases, the first round of PCR consisted in 3 PCR replicates per sample, targeting the DNA coding for the V4-V5 hypervariable region of 16S RNA ribosomal with degenerate primers (Parada et al. 2016) : 515F (GTGYCAGCMGCCGCGGTAA) and 926R (CCGYCAATTYMTTTRAGTTT). Two other primer pairs (18SV9 and ITS2) were amplified, added to libraries and sequenced together with 16S-V4V5. Each primer was flanked in its 5’-end by a nucleotide sequence used for indexing at a later step, according to a protocol proposed by Nag et al. (2017). At this stage, 2 additional PCR blanks were done with water instead of extracted DNA. Each 12,5 µl reaction mix contained 1 µl of DNA (∼10ng.µl^-1^), 0,25 µl of forward primer, 0,25 µl of reverse primer (10nM), 6,25µl of 2✕ Promega Green Master mix G2, 4,25µl of milliQ water. The PCR cycles consisted of of initial denaturing for 2 min at 94°C, followed by 30 cycles (denaturation 30 s at 94°C, hybridization 30 s at 51°C, elongation 45 s at 72 °C) and a final elongation during 5 min at 72°C. First PCR products were verified by electrophoresis on 1% agarose gel, re-amplified if negative until they were positive. Each PCR triplicate was pooled into one before the indexing PCR. Indexation PCR was realized in a 27.5 µl reaction mix containing 2 µl of first PCR products, 5 µl of reverse and forward index, 12,5µl of NEB Q5 2X mix and 8µl of milliQ water. This second PCR consisted of a initial denaturing for 30s at 98°C, followed by 30 cycles (denaturation 20s at 98°C, hybridization 20s at 60 °C, elongation 10s at 72°C) and final elongation 10s at 72°C. At this stage, one PCR blank was added with water instead of first PCR products. All indexed samples were pooled into a single low-bind tube and purified with magnetic beads (Nucleomag, Macherey Nagel, 1:1 ratio). Size range of final PCR products was verified by electrophoresis (Agilent BioAnalyzer, High-sensitivity), with an waited size peak around 420bp, then pooled in a final library, and sequenced on an Illumina MiSeq (one Miseq Reagent v3 kit 600 cycles and one nano MiSeq Reagent kit v2 kit 500 cycles for resequencing) in the Concarneau marine station (MNHN) to output demultiplexed fastq files.

### 2.4 Nanopore library

The same DNA extracts were processed in parallel for Nanopore 16S-FL sequencing, with the following 16S markers : V1-V9 regions (for bacteria, ∼1.45 kpb; Weisburg et al. 1991; 27F:AGAGTTTGATCMTGGCTCAG ; 1492R: TACGGYTACCTTGTTACGACTT). PCRs were performed in 3 small-volume replicates of 12,5 µl each, containing 6,25µl of LongAmp Taq 2✕ Master Mix (NEB), 4,25µl of milliQ water, 1 µl of DNA (∼10ng.µl^-1^), 0,25 µl of forward primer, 0,25 µl of reverse primer (10nM each). PCR cycles consisted of initial denaturing for 3 min at 94°C, followed by 30 cycles composed of denaturation for 30s at 94°C, hybridization for 30s at 51°C, and elongation for 45s at 65°C and final elongation for 10 min at 65°C. All first PCR products were verified by agarose gel electrophoresis, re-amplified if negative until they were positive, and positive triplicates were pooled into one before the indexation PCR. Concentrations were measured by the Qubit fluorometer (dsDNA BR kit) and brought back to a concentration of 1ng.µl^-1^. Indexation PCR was realized according to the Nanopore « PCR barcoding (96) amplicons (SQK-LSK109) » protocol. Indexed amplicons were pooled into one tube per primer/marker and purified with magnetic beads (Nucleomag Macherey Nagel, 1:0.8 ratio). Indexed and purified products were verified on agarose gel electrophoresis.

DNA concentration was measured by phospho-luminescence (Qubit), then diluted in order to have 1µg of DNA into 47µl of water. Final ligation of Nanopore sequencing adapters was done following the “SQK-LSK109 with EXP-PBC096” protocol. 16S V1-V9 library was sequenced on two R9.4.1 MinION flow cells (half of the samples + Ze for each). Flow cells were loaded on MinION Mk-1C and sequenced for approximately 48h, until no further sequencing reads could be collected above Q10 quality score. Fast5 files were basecalled and demultiplexed using Guppy 6.4.2 high-accuracy model on a local GPU (Nvidia Quadro K4000) and DNA sequence reads were output with >Q10 flag, as fastq files. For Illumina 16S-V4V5 and Nanopore 16S-FL, samples with less than 1500 reads were re-sequenced.

Sequence data are available in NCBI with BioProject accession number PRJNA985243.

### 2.5 Processing of raw reads

Fastq files from Illumina 16S-V4V5 were filtered with R package DADA2 v 1.16.0 (Callahan et al. 2016). Reads R1 and R2 were filtered using the *filterAndTrim* function (minLen=200, matchIDs=TRUE, maxN=0, maxEE=c(3,3)), then merged to unique sequences (ASVs) with at least 12 overlapping nucleotides between R1 and R2. Chimeric sequences were removed using the *removeBimeraDenovo* function. A matrix of 16S-V4V5 ASVs per sample was obtained and processed by Qiime2 tools (Hall et al. 2018), after 16S-V4V5 ASVs were extracted from fasta files containing sequences from other primers (18SV9 and ITS2, not presented here). Nanopore 16S-FL fastq sequences (>Q10) were filtered with Nanofilt (De Coster et al. 2018) : all reads shorter than 1.4 kpb and longer than 1.6 kpb for 16S V1-V9 were removed. *In-silico* V4V5 16S-V4V5 were extracted from mock sample 16S-FL with seqkit amplicon tool (Shen et al. 2016). Then, for all samples, ASVs with 97% of similarity were clustered into OTUs using the Vsearch (version 2023.7.0) tool, producing a table of OTU abundances from a table of ASV abundances. OTUs were taxonomically assigned with a trained Qiime2 classifier (confidence >0.75), inferring to the SILVA NR 99 reference database v138.1 (Quast et al. 2013), formatted for this specific marker.

### 2.6 Community structures analysis

Chloroplastic, mitochondrial and eukaryotic assignments, contaminants detected from blanks and singletons (OTUs with only 1 read in all samples) were removed from OTU tables. Tables of filtered OTU read abundances, OTU taxonomy and sample data were imported to make phyloseq objects in R, one for each marker (R package phyloseq, McMurdie and Holmes 2013).

The prokaryotic community structure of environmental samples depends tightly on the read number in each sample. The conventional rarefaction consists in randomly depleting reads in each sample, until all samples reach the number of reads of the poorest one (Simberloff 1972). This method is known to have major bias: non-reproducibility since reads are removed randomly, and alteration of community structures due to the random sorting of rare species (Coddington et al. 2009). In soil or sediment microbiotas, sample OTU richness depends strongly on sample size, therefore we opted for the rarefaction method developed by Chao and Jost (2012), consisting in comparing samples of equal completeness (equal *coverage*), not of equal size. “When samples are standardized by their coverage (a measure of sample completeness […]) instead of by their size, the estimated richnesses approximately satisfy a replication principle, which is an essential property for characterizing diversity” (Chao and Jost 2012). This coverage-based rarefaction was used by the function *phyloseq_coverage_raref* (R package metagMisc, Mikryukov 2019).

Since the 16S-V4V5 primers (16S-V4V5) amplified both bacteria and archaea, but 16S-FL primers (16SV1-V9) amplified bacteria only, archaeal taxa obtained by 16S-V4V5 were removed for the present analysis (deleted after rarefaction). Further analyses with archaeal taxa are however proposed in Supplementary Materials.

Analyses were carried out on filtered OTU tables after coverage-based rarefaction, except for Fig. 5, in which both rarefaction methods are shown. In sediment samples, core members were identified by their prevalence among all samples (≥50%, i.e. they were present in 50% or more of the samples, Fig. 5). For exploring dissimilarities between datasets, a Principal Coordinate Analysis (PCoA, from *phyloseq ordinate* function, equivalent to MDS - Metric Multidimensional Scaling) was performed on matrices of Bray-Curtis distances between communities. To identify the most contributing OTUs to the different parts of the communities, a Principal Component Analysis (PCA, from R package ade4, *dudi.pca* function) was performed on relative abundances. In order to assess the similarity of community structures described by both sequencing methods, a Procrustes analysis was carried out on their respective PCoA scores, with *procrustes* and *protest* functions (R package vegan). In parallel, a co-inertia analysis on PCA two first components was done, with *coinertia* and *RV.rtest* (999 permutations) from ade4. A Mantel permutation test was performed on two matrices of Bray-Curtis distances, for 16S-V4V5 and 16S-FL bacterial communities (Pearson method, 999 permutations), with vegan R package. Stochasticity was assessed in bacterial communities from each site, either from 16S-V4V5 and 16S-FL sequences with NST R package (Normalized Stochasticity Ratio in community assembly, Ning et al. 2019). Classification trees were used to characterize the genus and species contributing the most to the [site x (sea-land orientation] effect in each dataset by the R package randomForest (Liaw and Wiener 2002).

## 3 RESULTS

### 3.1 Mock Community

Mock community sequenced in 16S-V4V5 on Illumina was quite accurate, with all awaited bacterial genus correctly detected (*Bacillus, Limosilactobacillus [Lactobacillus]*, *Salmonella, Escherichia*, *Listeria, Enterococcus, Pseudomonas, Enterococcus* and *Staphylococcus*), and even the Eukaryota *Cryptococcus* by mitochondrial DNA. Noteworthy, (i) almost all genus were represented by a single OTU (97%), except *Escherichia* and *Limosilactobacillus* (2 species each) and (ii) 2 contaminating OTUs were detected (unknown *Bacteria* and *Exiguobacterium* sp., Fig. 2a-b).

**Figure 2.**
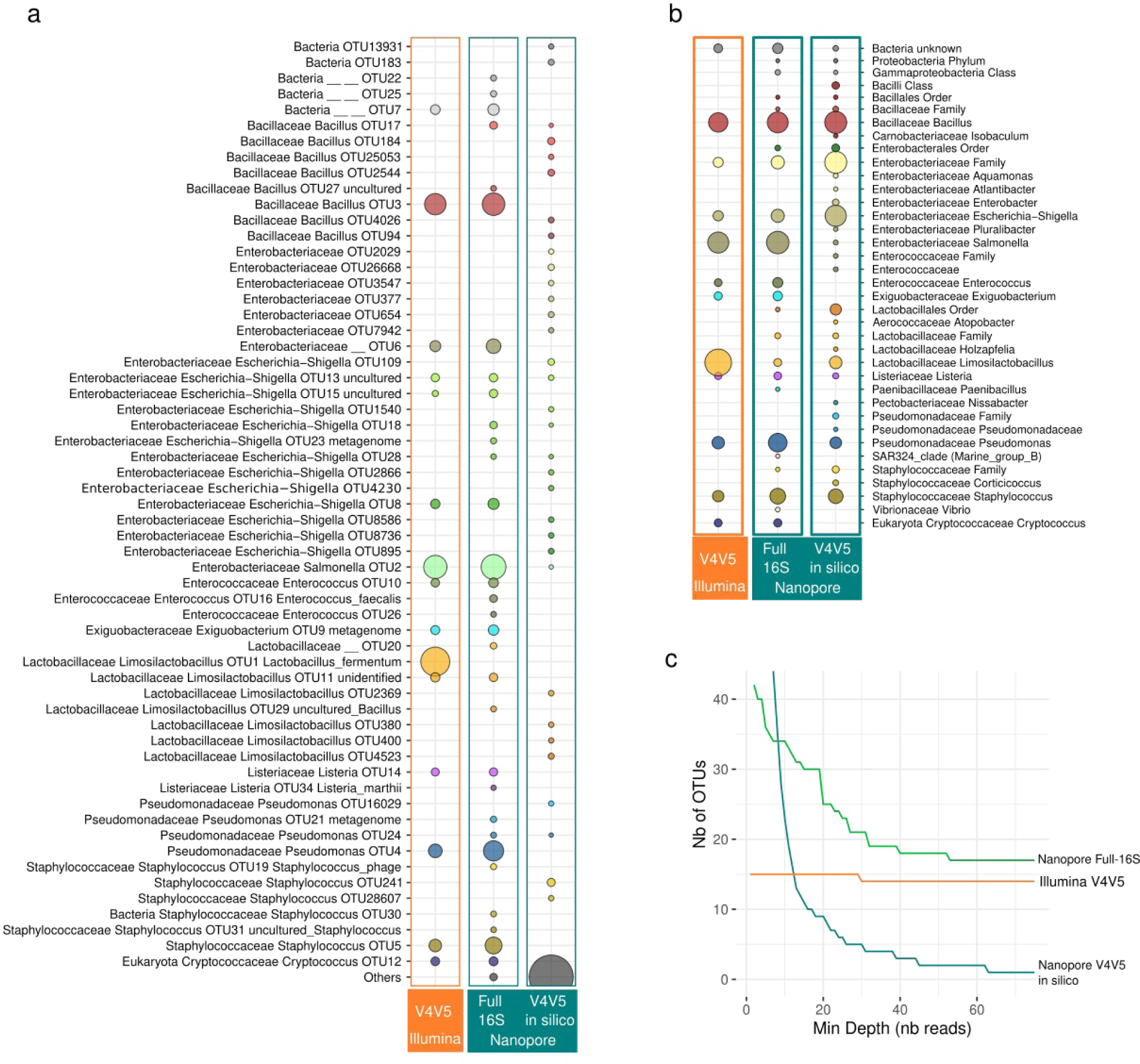
Relative abundances found in a mock community sequenced either by SR (16SV4V5 on Illumina), LR (full-16S on Nanopore), and in-silico SR (V4V5 extracted in silico from Nanopore LR) after singletons filtering and equal read rarefaction (9366 reads per sample) : (**a**) aggregated at species level (97%) ; (**b**) aggregated at genus level ; (**c**) evolution of the number of OTUs (97%) related to the minimum number of reads per OTU, depending on sequencing strategies.

The same mock community sequenced in 16S-FL on Nanopore found the very same species than those found by 16S-V4V5, except one of the two *Limosilactobacillus*. Additional species (37) belonging to 11 genus (Fig. 2a) were also found, albeit these latter only accounted for 9.8 % of the filtered reads for this community. The extraction *in silico* of the V4V5 domain from the 16S-FL did not detect the great majority of the species actually in the mock sample (Fig. 2a), but detected 9 genus over the 12 detected by 16S-V4V5, and only missed *Enterococcus* (Fig. 2b). When OTUs were filtered with an increasing minimal read depth (from 2 to 75, Fig. 2c), 16S-V4V5 and 16S-FL communities converged toward 14 and 17 species respectively, whereas the community described by *in silico* extracted V4V5 from 16S-FL collapsed completely.

### 3.2 Samples read coverage

After removing non-bacterial taxa and coverage-based rarefaction, the final mean abundance of Illumina 16S-V4V5 was mean 2609±828.1 reads per sample (min 1338, max 4991 reads, 749 OTUs), those of Nanopore 16S-FL was 5108±346.5 reads per sample (min 4451, max 6019 reads, 1495 OTUs). With conventional read rarefaction, for bacteria only, all samples were standardized at 1582 reads for both sequencers, resulting in a total of 570 (16S-V4V5) and 967 (16S-FL) bacterial OTUs, so in proportion 16S-FL counted 170% of the OTUs (species rank) detected by 16S-V4V5 (Table 1). In proportion with this rarefaction method, 16S-FL detected twice more species than 16S-V4V5. For the rest of this section, only results obtained by the coverage-based rarefaction method are presented.

**Table 1.** Statistics on bacterial taxa detected by 16S-V4V5 and 16S-FL primer pairs, shared or unshared taxa between primer pairs, by taxonomic rank, percentage of taxa or reads assigned. Here only the bacterial dataset was coverage-based rarefied and singletons filtered. *among taxa assigned at this rank (non-assigned are not counted). In the bacterial communities analyzed, 84.7 and 98.8% of 16S-V4V5 were assigned to the same species and genus, respectively, than those detected by 16S-FL (in orange). Conversely, 68.5 and 89.6% of 16S-FL were assigned to the same species and genus, respectively, than those detected by 16S-V4V5 (in blue).

| Primers / Sequencer | Taxa / Reads | Phylum | Class | Order | Family | Genus | Species |
| --- | --- | --- | --- | --- | --- | --- | --- |
| **16S-V4V5**<br>135 692 reads<br>Illumina<br>(515F + 926R) | **Taxa detected** | **45** | **106** | **209** | **309** | **448** | **749** |
|  | % taxa assigned | 97.8% | 92.5% | 91.9% | 88.0% | 82.4% | 65.3% |
|  | % reads assigned | 99.6% | 99.6% | 98.3% | 98.0% | 94.0% | 54.0% |
|  | **Taxa exclusive** | **2** | **7** | **15** | **24** | **55** | **227** |
|  | % taxa assigned | 100% | 100% | 93.3% | 87.5% | 81.8% | 80.6% |
|  | % reads assigned | 100% | 100% | 97.1% | 83.9% | 84.5% | 85.8% |
|  | % taxa shared with 16S-FL<br>% of the total 16S-V4V5 reads with same assignment than 16S-FL* | 95.6%<br>99.99% | 93.4%<br>99.97% | 92.8%<br>99.7% | 92.2%<br>99.7% | 87.7%<br>98.8% | 69.7%<br>84.7% |
| **16S-FL**<br>265 650 reads<br>Nanopore<br>(27F + 1492R) | **Taxa detected** | **54** | **140** | **316** | **483** | **785** | **1495** |
|  | % taxa assigned | 98.1% | 89.3% | 89.2% | 85.9% | 80.9% | 64.6% |
|  | % reads assigned | 88.3% | 86.0% | 74.9% | 73.3% | 67.8% | 46.9% |
|  | **Taxa exclusive** | **11** | **41** | **122** | **198** | **392** | **973** |
|  | % taxa assigned | 100% | 82.9% | 85.2% | 82.8% | 79.3% | 67.8% |
|  | % reads assigned | 100% | 70.5% | 87.2% | 83.7% | 81.8% | 67.4% |
|  | % taxa shared with 16S-V4V5<br>% of the total 16S-FL reads with same assignment than 16S-V4V5* | 79.6%<br>99.7% | 70.7%<br>99.3% | 61.4%<br>95.7% | 59.0%<br>94.7% | 50.1%<br>89.6% | 34.9%<br>68.5% |
| Shared taxa | **Shared taxa** | **43** | **99** | **194** | **285** | **393** | **522** |
|  | % taxa shared | 76.8% | 67.3% | 58.6% | 56.2% | 46.8% | 30.3% |
|  | % taxa assigned among shared | 97.6% | 91.9% | 91.8% | 88.1% | 82.4% | 58.6% |

### 3.3 Variations in community structures

Community composition and multivariate analyses showed that both technologies detected a marked difference between bacterial communities from Babin and Rivière salée sites, but also their fine-scale orientation, from sea- to land-oriented samples. Communities sequenced by 16S-V4V5 and 16S-FL described the same global patterns, i.e. a preponderance of Pirellulales (Planctomycetota) in Rivière salée, of Pseudomonadales (Gammaproteobacteria) and Bacteroidales (Bacteroidota) in Babin, separating clearly the two sites in ordination (Fig. 3a-b).

**Figure 3.**
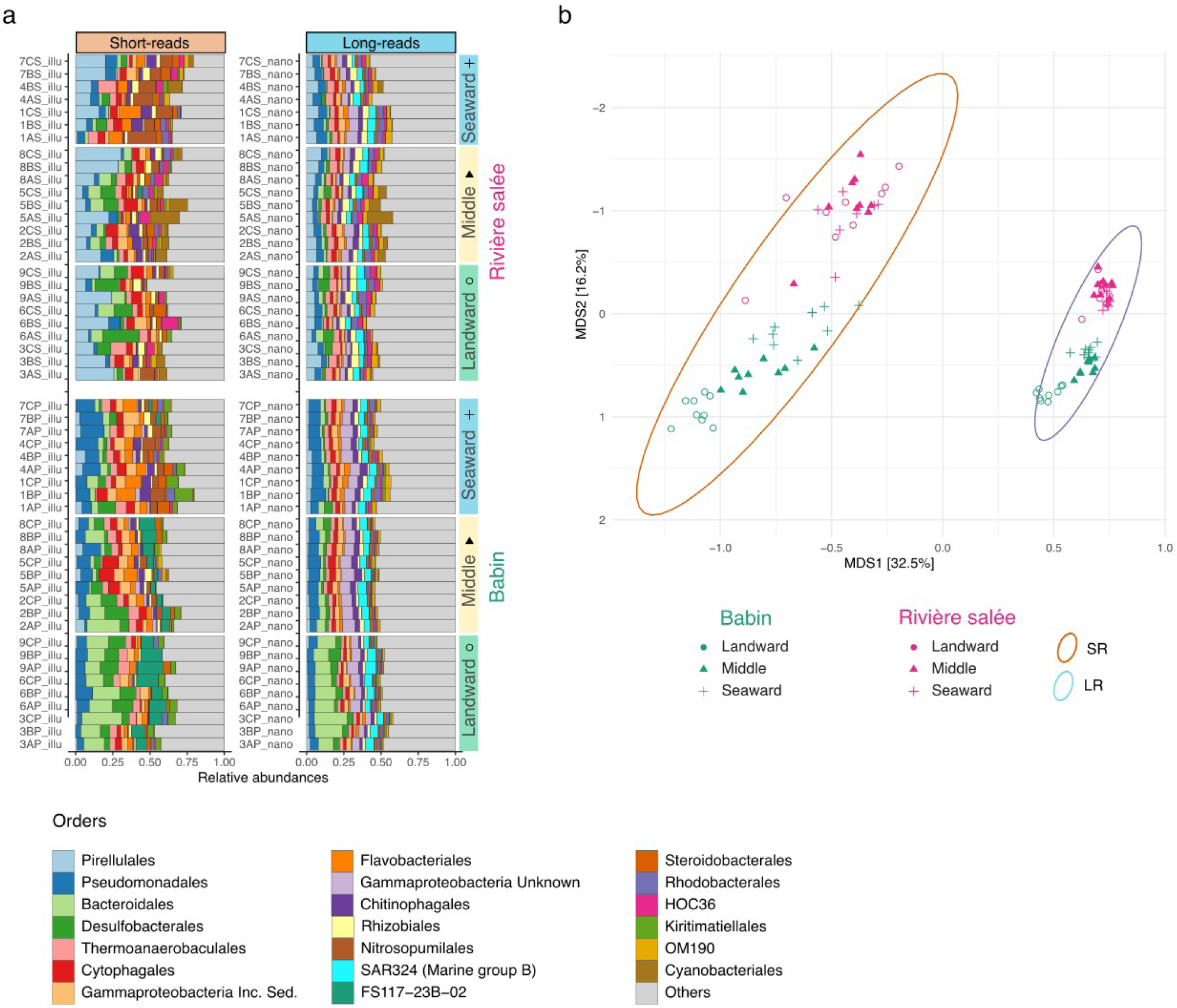
(**a**) Top-20 bacterial orders in samples for both sequencing devices, ranked by their overall relative abundances in samples ; (**b**) biplot of sample scores from a nMDS on abundances of bacterial OTUs agglomerated at genus level, for both sequencing devices (stress=14.1%) ; for this common ordination, shared OTUs were named differently between SR and LR on purpose, in order to separate the two datasets for a better visualization. Number of reads per sample was rarefied with the coverage-based method (Chao and Jost 2012).

Babin showed the most structured community along the tidal gradient, with the presence of Pseudomonadales in seaward samples and of Bacteroidales (Bacteroidota) in landward samples. Biological replicates were relatively close to each other in the PCoAs (Fig. 4a-b), but Bray-Curtis dissimilarity indexes of communities within replicates were always higher for 16S-V4V5 than for 16S-FL, either for Babin or Rivière salée (Fig. 4c, anova p<0.001).

**Figure 4.**
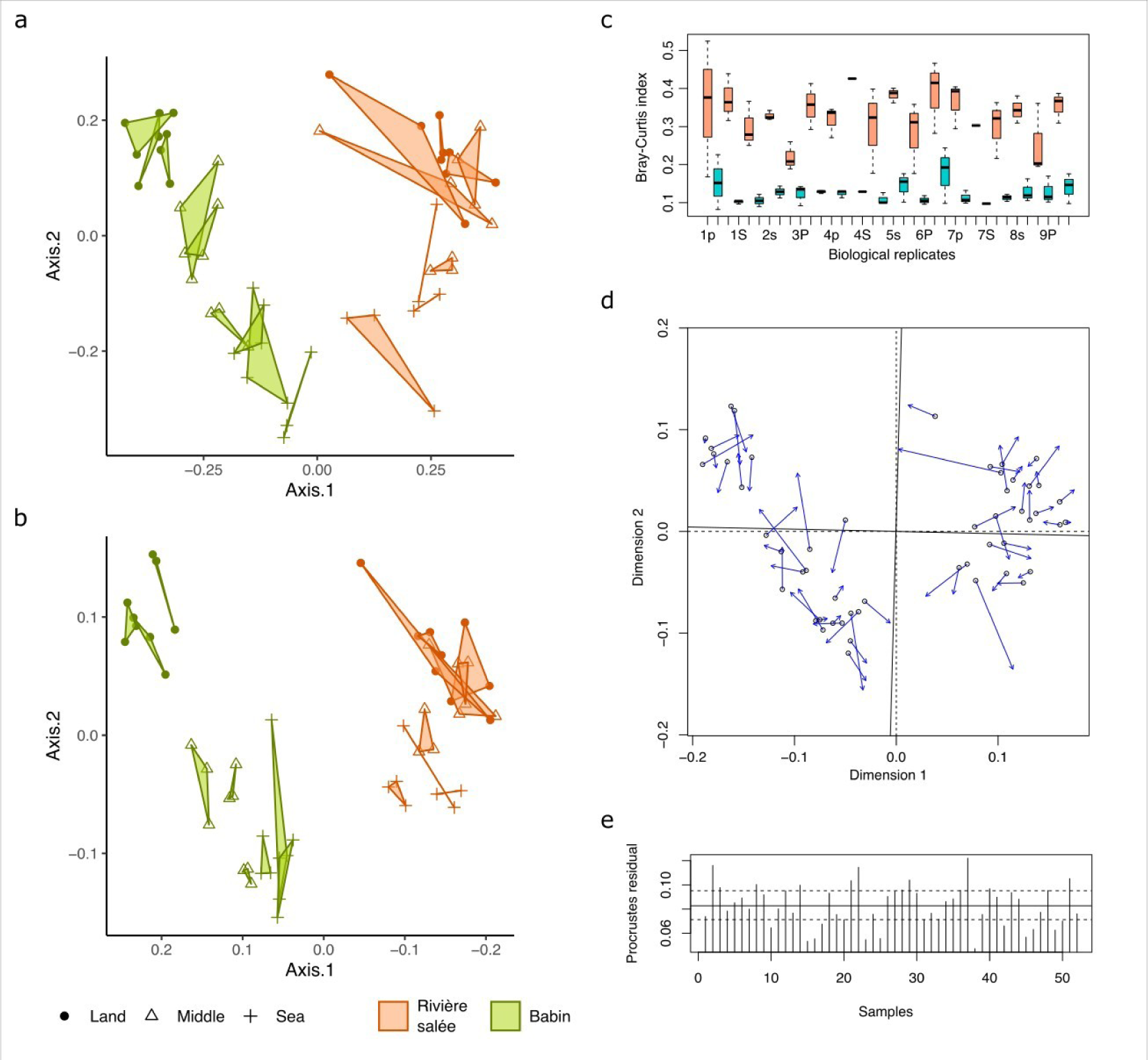
(**a-b**) PCoA on coverage-based rarefied abundances of bacterial communities at species level, (**a**) sequenced by SR, showing biological replicates (polygons) ; (**b**) sequenced by LR ; (**c**) Dispersion of Bray-Curtis dissimilarity index within biological replicates, salmon boxplots for SR, cyan for LR ; thick horizontal lines : mean ; box plots : 75% range ; whiskers : 95% range ; dots: outliers ; (**d**) Procrustes analysis of the 2 first components of both PCoAs (presented in a-b), showing the degree of matching between the two ordinations ; empty dots show the position of the samples in the LR ordination and arrows point to their positions in the SR ordination ; the plot also shows the rotations between the axis (solid *vs*. dashed), necessary to make ordinations match as closely as possible ; (**e**) residuals for each sample between the ordinations (this time, on the 20 first axis); the horizontal lines, from bottom to top, are the 25% (dashed), 50% (solid), and 75% (dashed) quantiles of the residuals.

**Figure 5.**
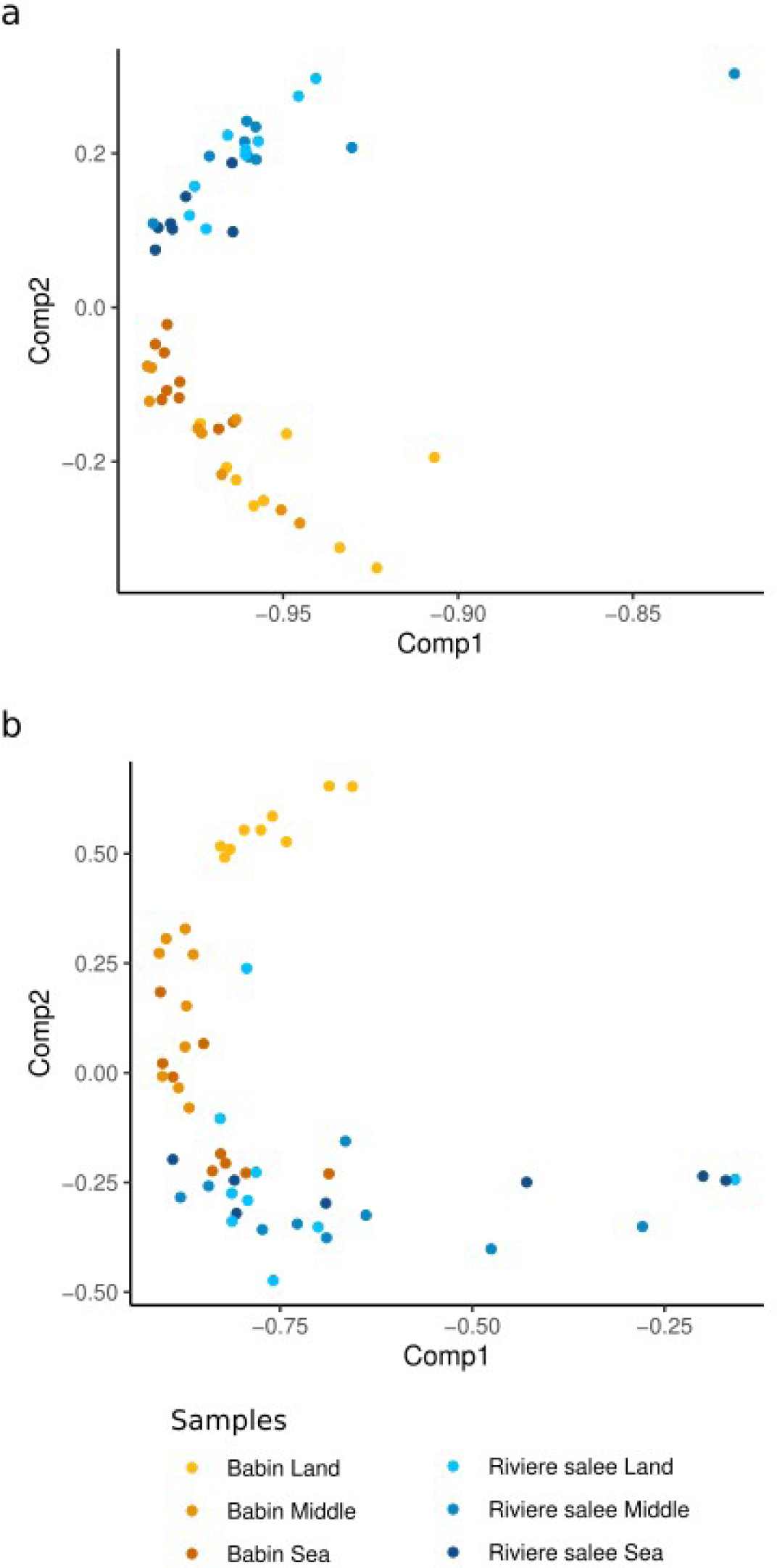
PCA on (**a**) species OTUs common to both 16S-V4V5 and 16S-FL (N=961), with samples colored according to sites, (**b**) species OTUs only detected by 16S-FL (N=634).

**Figure 6.**
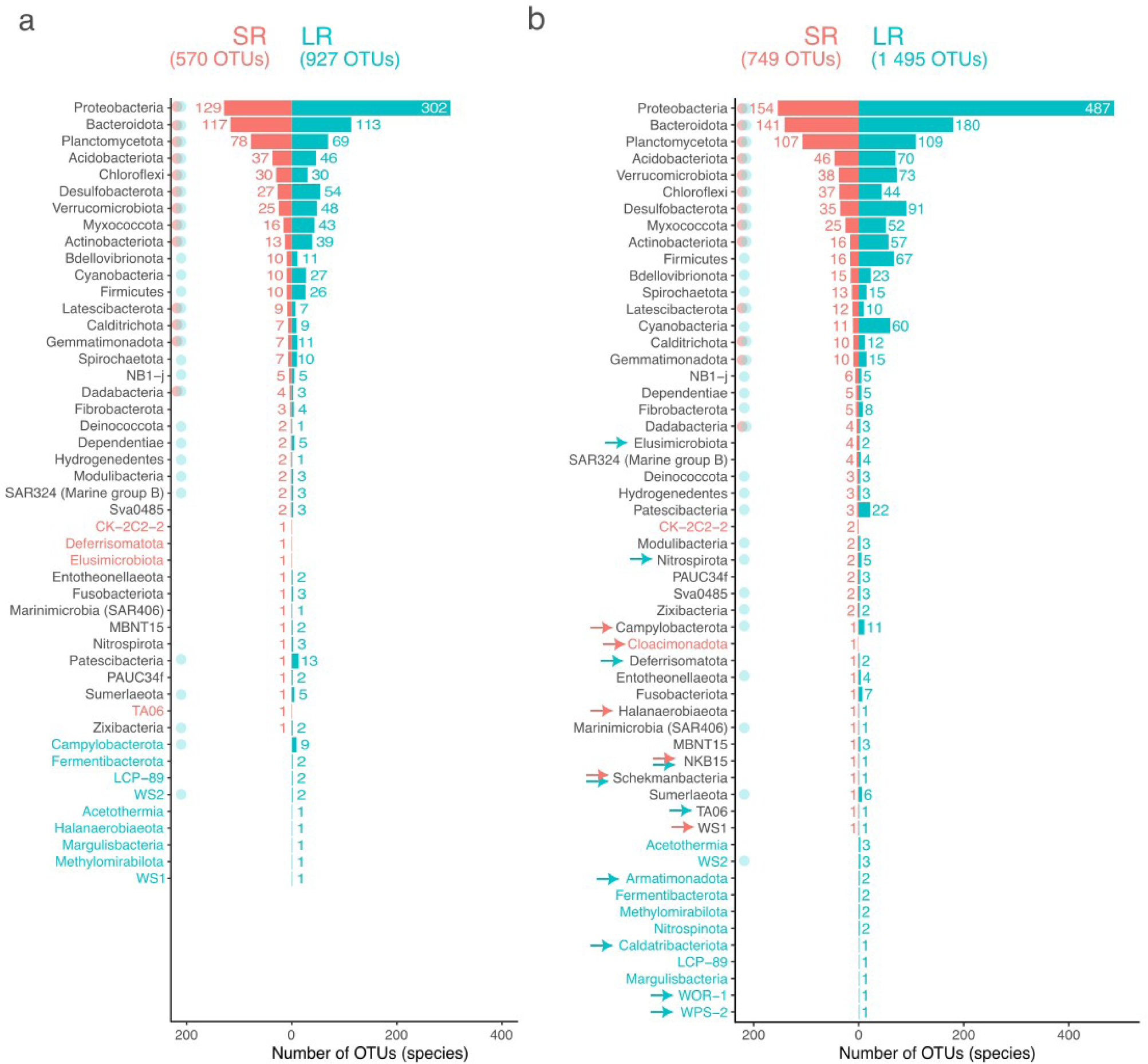
Number of OTUs (97% similarity, singleton-filtered) for each prokaryotic phylum in environmental samples analyzed here (bacteria only), depending on the sequencing strategy : (**a**) with conventional equal-rarefaction (1582 reads for all samples of both strategies, see Methods section for comments on inner bias) ; (**b**) with coverage-based read rarefaction (see Results section for details). Phylum names in red or blue were detected only by SR or only by LR, respectively. Red or blue dots indicate core-phyla, i.e. phyla with a minimum prevalence of 50% in the respective datasets. Red or blue arrows indicate phyla that were not detected with read equal-rarefaction, for SR or LR respectively.

The Procrustes analysis of the two first axes of multivariates showed a significantly strong similarity between structures drawn by 16S-V4V5 and 16S-FL (Fig. 4d-e, correlation 0.793, p<0.001), confirmed by a co-inertia analysis on PCA’s two first axes (p<0.001). The Mantel test indicated a significant correlation coefficient of 0.7248 (p<0.001) between the Bray-Curtis dissimilarity matrices obtained from 16S-FL and 16S-V4V5 communities using OTUs at species taxonomic rank (i.e. with or without a species level taxonomic assignment).

Stochasticity of bacterial communities calculated based on the normalized stochasticity ratio (NST) was high for all groups and sequencing strategies : 84.6% / 85.4% for Babin / Rivière salée sites from bacterial species common to both 16S-FL on Nanopore and 16S-V4V5 on Illumina ; 80.0% / 87.8% for the same sites from bacterial species exclusive to 16S-FL on Nanopore.

In order to point out the similarity of taxa contributing to the [site ✕ (sea-land orientation] effect, classification trees were made by a random forest approach on the 393 genus and 285 families shared between 16S-V4V5 and 16S-FL. Models found 48% of similarity among the top-100 contributing genus and 63% among the top-100 contributing families between sequencers. However, taxa contributing in the same way to the [site ✕ (sea-land orientation] effect were scarce (Fig. S1, Table S1). Archaean communities described with 16S-FL specific primers followed roughly the structure obtained with bacteria (Fig. S2).

### 3.4 Phylogenetic diversity

Over the 56 bacterial phyla detected in total, 54 were detected by 16S-FL and 45 by 16S-V4V5 (Table 1). At high taxonomic levels, 16S-V4V5 and 16S-FL were approximately ⅘ alike for phyla, 16S-FL detected 11 exclusive phyla over a total of 54 for this primer (20% of exclusive among those detected by 16S-FL), when 16S-V4V5 only had 2 (4.4%). However, 16S-FL-exclusive subcommunity represented only 0.2% of reads in the full community (Fig. S3). The 11 phyla only detected by 16S-FL were *Acetothermia*, *WS2*, *LCP-89*, *WOR-1*, *Armatimonadota*, *Margulisbacteria*, *Nitrospinota*, *Fermentibacterota*, *Methylomirabilota*, *Caldatribacteriota*, *WPS-2*, whereas the only 2 detected by 16S-V4V5 were *Cloacimonadota* and *CK-2C2-2,* with a coverage-based rarefaction (Fig. 5). At lower taxonomic levels, 92.2% and 87.7% of respectively the family and bacterial genus detected by 16S-V4V5 were detected by 16S-FL. 16S-FL detected twice more species than 16S- V4V5, with only 34.9% of the species and 50.1% of the genus detected shared with 16S-V4V5. The trend that 16S-FL detected almost all 16S-V4V5 taxa *plus* a certain number of 16S-FL original taxa decreased with lowering taxonomic ranks (Fig. S4).

All the 54 16S-FL-detected phyla were more diversified based on 16S-FL, but four : *NB1-j*, *SAR324*, *Dadabacteria* and *Hydrogendentes*. The most diversified phylum, the *Proteobacteria*, presented more than 4 times more species with 16S-FL than with 16S-V4V5. Overall, communities described by the two primer-sets were phylogenetically very similar when considering shared taxa at the family and genus level (92.2% and 87.7% of taxa similarity for 16S-V4V5 *vs.* 16S-FL, respectively).

11.7% of the 16S-FL (bacterial 16S) were unassigned at the phylum level, versus 0.36% for 16S- V4V5. 53.1% of the 16S-FL unassigned at the species level (35.4% of total 16S-FL bacterial OTUs), versus 46.0% for 16S-V4V5 (34.7% of total 16S-V4V5 bacterial OTUs, Table 1). For shared genera, the unassigned reads were much lower for 16S-V4V5 (5.8%) than 16S-FL (35.5%). All core-phyla detected by 16S-V4V5 were also parts of core-phyla detected by 16S-FL, whatever the rarefaction method used (Fig. 5).

## 4 DISCUSSION

In this study, bacteria rRNA was amplified on their gene 16S-V4V5 and 16S-FL primers from the same extractions of environmental samples, then sequenced on Illumina and Nanopore respectively, and assigned on the same database of reference sequences.

The error rate of Nanopore (around 6 %, Tyler et al. 2018) did prevent its use for in-silico-extracted 16S-V4V5 in the present study, but the read lengthening allowed to catch up a sufficient accuracy and to describe the bacterial communities from marine sediment samples in consistency with Illumina 16S-V4V5, either in the coarse structure (site effect) and fine structure (sea-land orientation), with nonetheless a couple of constant differences, already noticed with the assessment of Katiraei et al. (2022) comparing 16SV4 *vs*. 16S-FL sequencing on Illumina and PacBio : (i) communities described by 16S-FL were more species-diversified than those described by 16S- V4V5, which is known to be at least partially due to differences in primer pairs ; (ii) abundances of OTUs based on 16S-FL were slightly less variable within biological replicates than those based on 16S-V4V5. Thus, the present work suggests that 16S-FL can be used for metabarcoding bacteria communities from environmental samples on a Nanopore sequencing device.

### 4.1 Higher error rate of Nanopore partially caught up by read lengthening

Katiraei et al. (2022) sequenced 16S-FL amplicons on a PacBio system, and extracted afterward in silico the 16SV4 fragments. In-silico-extracted V4 dataset had approximately half of the read count per sample, compared to those of the 16S-FL PacBio dataset, indicating that a significant proportion of the taxa that were identified by 16S-FL were not detected by extracting the V4-region from the same initial sequences. In this way, the length of the 16S fragment can modify the taxonomic assignment, a longer fragment increasing the diversity of taxa assigned, albeit not figuring if they were true taxa or not. Our study confirmed that there were much more taxa detected by 16S-FL than by 16S-V4V5, but also that a certain proportion of taxa sequenced with 16S-V4V5 were not detected with 16S-FL dataset (30.3% of the species and 12.3% of the genus, Table 1). However, this observation was much tempered when the proportion of reads involved in these non-detected taxa was considered, concerning 15.3% of the reads for species and only 1.2% for genus.

When considering non-shared taxa, the present study illustrated the assignation power of a longer bacterial 16S rRNA, compared to a restricted 16S V-region, incidentally acknowledged to have the most appropriate cover for bacteria among 16S-V4V5 primers (Parada et al. 2016; Walters et al. 2016; Willis et al. 2019). Taxa assignment rates were lower at species level whatever the read length, probably due to the incomplete databases that are constantly being updated, or pseudogenes and intra-genome 16S polymorphism (Pei et al. 2010; Větrovský and Baldrian 2013), impossible to evaluate with our approach.

### 4.2 Phylogenetic and ecological patterns conserved

Coarse and fine spatial structures were overall significantly similar, since the site effect and the sea-land orientation were conserved in ordinations.

Differences in abundances for the same taxa were obvious in the structure of mock communities, i.e. coming from the same DNA extraction but followed by separate amplification on different primers, different library preparation and sequencing. This discrepancy is typical and outlines the semi-quantitative trait of any microbial HTS sequencing. However, all qualitative elements (beta-diversity) of mocks were preserved, allowing us to extend this observation to communities described from environmental samples processed with the same workflow as for the mock. This assumption of a correct taxa detection in spite of abundance discrepancies may explain differences observed in top-20 bacterial taxa influencing structures (Fig. 3a), and is reinforced by the relative orientation of samples, preserved between the two sequencing workflows on the same ordination (Fig. 3b).

On the other hand, it is noteworthy that 16S-FL communities contained twice more OTUs than 16S- V4V5 ones and this did not change the overall structure of ordinations, providing evidence that core-communities in both sequencing strategies were congruent and that additional taxa detected by 16S-FL did not significantly changed this ordination. In another perspective, 16S-V4V5 Illumina’s communities, albeit reduced, were sufficient and contained the smallest share of taxa needed to correctly describe the assemblages at play.

Our study on marine sediment samples could not provide evidence that 16S-FL improved the taxonomic assignment, as it was done with human gut microbial communities (Jeong et al. 2021; Matsuo et al. 2021). However, if genus level is considered as the maximum resolution of 16S sequencing for a correct taxonomic assignment, a proper primers comparison should start from genus toward higher levels. The fact that read assignment was always lower for 16S-FL-exclusive taxa probably reflects more the fact that mangrove sediments contain a high diversity of uncultivated microbes with presently unavailable 16S-FL in reference databases, than a lower sequencing accuracy of Nanopore (and therefore a plausible sequencing-platform effect).

## ACKNOWLEDGEMENTS

This work was lead by the GT DCE Mangroves (Groupe de Travail Directive Cadre sur l’Eau Mangroves), founded by the OFB (Office Français de la Biodiversité, Olivier Monnier), lead by the MNHN (Muséum National d’Histoire Naturelle, GD). Lab work was achieved by AL during her Master practice, with BOREA financial support. Thanks to our colleague Cédric Hubas (BOREA), for discussion and statistical advice. Thanks to associate-editor and reviewers of PCI microbiology for their pertinent suggestions.

## DATA ACCESSIBILITY

All data presented and scripts in this manuscript are available from the GitHub repository : https://github.com/tonyrobinet/nanopore_metabarcoding. Raw sequences (fastq format) are available in NCBI with BioProject accession number PRJNA985243.

## AUTHOR CONTRIBUTIONS

G.D. and T.R. designed the study and seeked for funds ; G.D. conducted the fieldwork and collected samples; T.R. designed the lab protocols, with support of A.L. ; A.L. performed the lab work, with the supervision of M.G. and T.R. ; T.R. and A.L. performed the statistical analysis ; T.R. wrote the manuscript, corrected by A.L. and G.D.

## SUPPLEMENTARY MATERIAL

Available at https://github.com/tonyrobinet/nanopore_metabarcoding

Figure S1. Contribution of Mean Decrease Gini coefficient (MDG) of common genus (a) and common families (b) sequenced by Illumina and Nanopore, for [site+(sea-land orientation)] predictors (see details in Table S2).

Figure S2. (a-b) Archaean taxa (genus level) contributing to structuring the communities in samples sequenced by Nanopore (rarefied at 5500 reads per sample, 97% OTUs with a minimum coverage of 50 reads) : (a) PCA on relative abundances, (b) iris plot of the relative abundances for taxa the most contributing to the PCA in (a). (c-d) same for bacterial taxa (genus level), sequenced by Nanopore.

Figure S3. Mean relative abundance of Nanopore phyla ; exclusive Nanopore phyla are in red.

Figure S4. (a) Venn diagrams showing the proportions of bacterial taxa shared and unshared between both primer pairs, at each phylogenetic rank (numbers in the discs refer to the numbers of taxa of the portion of the disc it is written on) ; (b) On left axis, proportion of bacterial taxa shared between both primer pairs (lines with dots) and proportion of reads assigned to taxa shared between both primer pairs (lines with stars), number of shared and unshared taxa (bars, right axis), at each phylogenetic rank ; more details in Table 1.

Table S1. Archaea detected by 16S-V4V5 were mentioned but the read coverage by sample was much lower than those for archaeal 16S-FL.

Table S2. Bacterial genus contributing the most importantly to the site effect, after a random forest analysis on Illumina and Nanopore datasets. In green : OTUs common to both datasets. MDG : mean decrease in Gini coefficient, a measure of how each variable contributes to the homogeneity of the nodes and leaves in the resulting random forest ; the higher the value of MDG score, the higher the importance of the variable in the model.

