## Supplementary materials for "Evaluation of a nanopore sequencing strategy on bacterial communities from marine sediments"

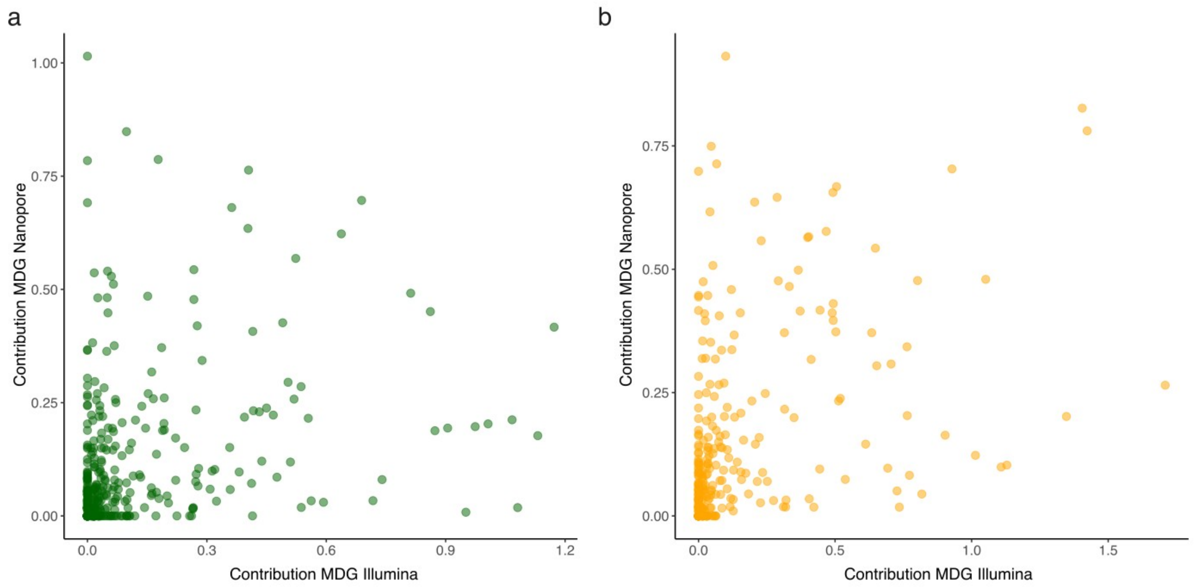


**Figure S1**. Contribution of Mean Decrease Gini coefficient (MDG) of common genus (**a**) and common families (**b**) sequenced by Illumina and Nanopore, for [site+(sea-land orientation)] predictors (see details in Table S2). Mean Decrease Gini is a measure of how each variable contributes to the homogeneity of the nodes and leaves in the resulting random forest (see Methods for details) ; the higher the value of MDG score, the higher the importance of the variable in the model.

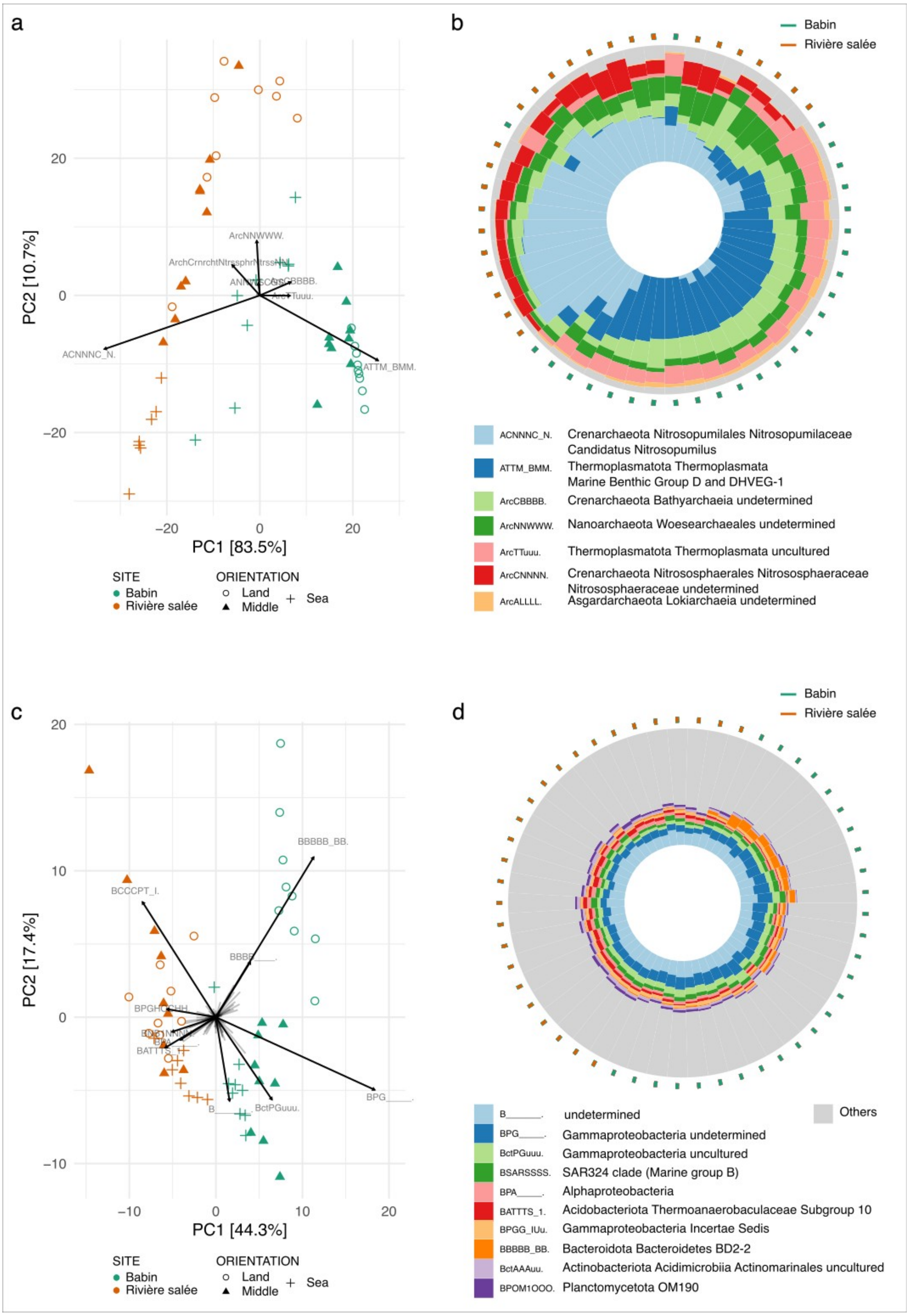


**Figure S2**. (**a-b**) Archaean taxa (genus level) contributing to structuring the communities in samples sequenced by Nanopore (rarefied at 5500 reads per sample, 97% OTUs with a minimum coverage of 50 reads) : (**a**) PCA on relative abundances, (**b**) iris plot of the relative abundances for taxa the most contributing to the PCA in (a). (**c-d**) same for bacterial taxa (genus level), sequenced by Nanopore.

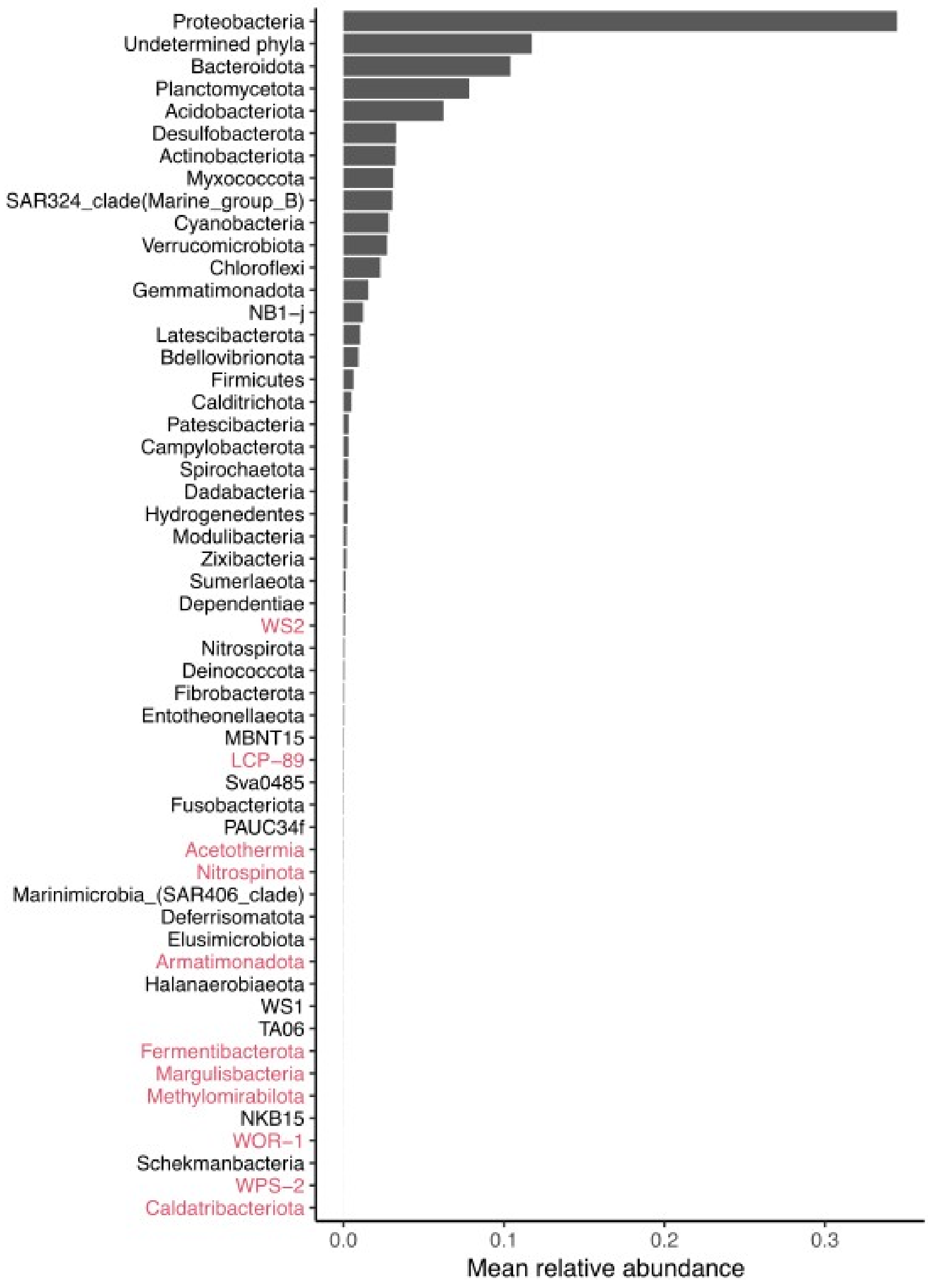


**Figure S3**. Mean relative abundance of Nanopore phyla ; exclusive Nanopore phyla are in red.

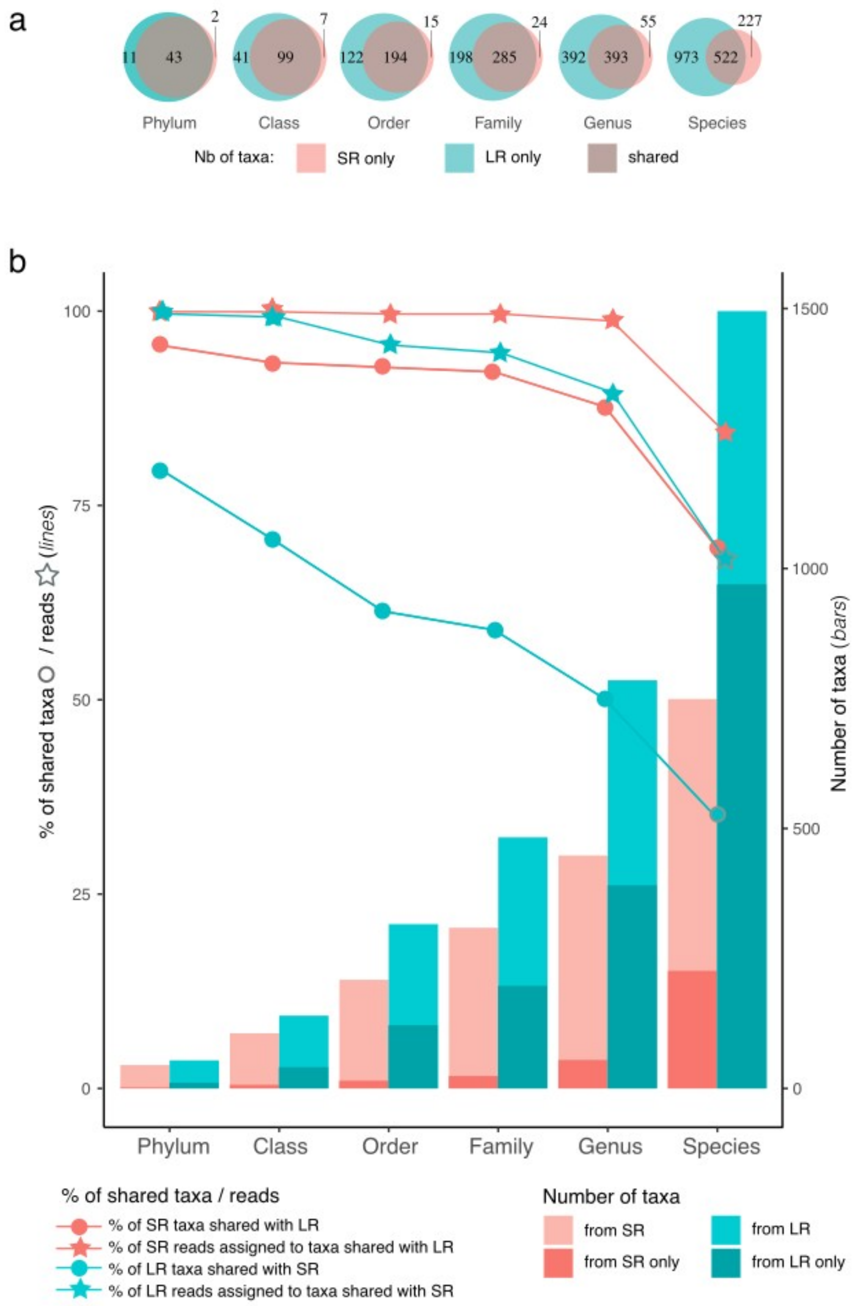


**Figure S4**. (**a**) Venn diagrams showing the proportions of bacterial taxa shared and unshared between both primer pairs, at each phylogenetic rank (numbers in the discs refer to the numbers of taxa of the portion of the disc it is written on) ; (**b**) On left axis, proportion of bacterial taxa shared between both primer pairs (lines with dots) and proportion of reads assigned to taxa shared between both primer pairs (lines with stars), number of shared and unshared taxa (bars, right axis), at each phylogenetic rank ; more details in Table 1.

| Kingdom | sequencer (primers) | Taxa / *reads* | Phylum | Class | Order | Family | Genus | Species |
|---|---|---|---|---|---|---|---|---|
| Archaea | Illumina (515F + 926R) | detected (% assigned) | 6 (83.3%) | 7 (85.7%) | 10 (90.0%) | 14 (85.7%) | 15 (86.7%) | 31 (71.0%) |
| | Nanopore (SSU1Ar F + SSU1000Ar) | detected (% assigned) | 11 (90.9%) | 23 (78.3%) | 34 (79.4%) | 50 (80.0%) | 83 (80.7%) | 171 (67.8%) |
| | shared | | 6 | 7 | 10 | 13 | 14 | 24 |
| | % of shared taxa for Illumina | | 100% | 100% | 100% | 92.9% | 93.3% | 77.4% |
| | % of shared taxa for Nanopore | | 54.6% | 30.4% | 29.4% | 26.0% | 16.9% | 14.0% |

**Table S1**. Archaea detected by Illumina were mentioned but the read coverage by sample was much lower than those for Nanopore Archaea.

| OTU# | Phylum | Class | Order | Family | Genus | MDG |
|---|---|---|---|---|---|---|
| | | | **Illumina** | | | |
| OTU499 | Dadabacteria | Dadabacteriia | Dadabacteriales | __ | __ | 2,26 |
| OTU295 | Planctomycetota | Planctomycetes | Pirellulales | Pirellulaceae | __ | 0,81 |
| OTU293 | | | | | *Blastopirellula* | 0,99 |
| OTU277 | | | | | *Pirellula* | 0,44 |
| OTU209 | Proteobacteria | Alphaproteobacteria | Rhizobiales | Rhizobiales inc. sed. | *Bauldia* | 0,40 |
| OTU161 | | Gammaproteobacteria | Ectothiorhodospirales | Ectothiorhodospiraceae | *Thiogranum* | 0,53 |
| OTU134 | | | HOC36 | HOC36 | *HOC36* | 1,13 |
| OTU111 | | | Pseudomonadales | Nitrincolaceae | *Marinobacterium* | 1,10 |
| OTU92 | | | Thiomicrospirales | Thiomicrospiraceae | *endosymbionts* | 2,48 |
| OTU107 | | | Pseudomonadales | Pseudohongiellaceae | *Pseudohongiella* | 0,50 |
| OTU118 | | | | Halieaceae | *Parahaliea* | 0,37 |
| OTU111 | | | | Nitrincolaceae | *Marinobacterium* | 1,10 |
| OTU139 | | | Gammaproteobacteria inc. sd. | Unknown_Family | *uncultured* | 0,39 |
| OTU190 | | | uncultured | uncultured | *uncultured* | 0,48 |
| OTU503 | Cyanobacteria | Cyanobacteriia | Phormidesmiales | Nodosilineaceae | *MBIC10086* | 0,60 |
| OTU52 | Verrucomicrobiota | Kiritimatiellae | Kiritimatiellales | Kiritimatiellaceae | *R76-B128* | 0,32 |
| OTU528 | Chloroflexi | Dehalococcoidia | FS117-23B-02 | FS117-23B-02 | *FS117-23B-02* | 0,60 |
| OTU637 | Bacteroidota | Bacteroidia | Flavobacteriales | Flavobacteriaceae | *Hoppeia* | 1,19 |

| | | | | | | |
|---|---|---|---|---|---|---|
| OTU643 | | | | | *Actibacter* | 0,36 |
| OTU703 | | | Bacteroidales | SB-5 | *SB-5* | 0,53 |
| OTU709 | | | | Prolixibacteraceae | *Draconibacterium* | 0,55 |
| | | | **Nanopore** | | | |
| OTU_b563 | Dadabacteria | Dadabacteriia | Dadabacteriales | __ | __ | 0,81 |
| OTU_b120 | Planctomycetota | Planctomycetes | Pirellulales | Pirellulaceae | __ | 0,73 |
| OTU_b118 | | | | | *Blastopirellula* | 0,39 |
| OTU_b84 | | | | | *Pir4_lineage* | 0,81 |
| OTU_b221 | Proteobacteria | Alphaproteobacteria | Rhizobiales | Rhizobiales inc. sed. | *Bauldia* | 0,75 |
| OTU_b1395 | | | Defluviicoccales | Defluviicoccaceae | *Defluviicoccus* | 1,14 |
| OTU_b18 | | Gammaproteobacteria | __ | __ | __ | 0,94 |
| OTU_b420 | | | Ectothiorhodospirales | Ectothiorhodospiraceae | *uncultured* | 0,42 |
| OTU_b157 | | | | Thioalkalispiraceae | *Thiohalophilus* | 0,86 |
| OTU_b98 | | | HOC36 | HOC36 | *HOC36* | 0,65 |
| OTU_b454 | | | Pseudomonadales | Nitrincolaceae | *Marinobacterium* | 0,85 |
| OTU_b1960 | | | | Alcanivoracaceae | *Ketobacter* | 0,81 |
| OTU_b355 | | | Thiomicrospirales | Thiomicrospiraceae | *endosymbionts* | 0,51 |
| OTU_b1138 | | | Chromatiales | Sedimenticolaceae | *Sedimenticola* | 0,85 |
| OTU_b365 | | | pItb-vmat-80 | pItb-vmat-80 | *pItb-vmat-80* | 1,47 |
| OTU_b202 | NB1-j | __ | __ | __ | __ | 0,77 |
| OTU_b502 | Acidobacteriota | Vicinamibacteria | Subgroup_9 | __ | __ | 0,77 |
| OTU_b115 | | Subgroup_22 | __ | __ | __ | 0,48 |
| OTU_b744 | Actinobacteriota | Actinobacteria | Corynebacteriales | Mycobacteriaceae | *Mycobacterium* | 0,48 |
| OTU_b862 | | Thermoleophilia | Solirubrobacterales | 67-14 | __ | 0,94 |

**Table S2**. Bacterial genus contributing the most importantly to the site effect, after a random forest analysis on Illumina and Nanopore datasets. In green : OTUs common to both datasets. MDG : mean decrease in Gini coefficient, a measure of how each variable contributes to the homogeneity of the nodes and leaves in the resulting random forest ; the higher the value of MDG score, the higher the importance of the variable in the model.
